# Oxytocin receptor dysfunction during critical neurodevelopment programs lasting pain hypersensitivity and sex-specific cognitive deficits

**DOI:** 10.64898/2026.07.04.736474

**Authors:** Hannah Illouz, Edith Tanche, Ophélie Schaack, Vincent Lelievre, Pierrick Poisbeau

## Abstract

Early life stress (ELS), modeled in rodents through neonatal maternal separation (NMS), induces lasting behavioral and molecular alterations including pain hypersensitivity, anxiety-like behaviors, and cognitive deficits. While NMS disrupts the oxytocinergic system, the specific contribution of oxytocin receptor (OTR) dysfunction during critical neurodevelopmental periods remains unclear. Here, we investigated whether neonatal OTR blockade alone could recapitulate key features of the NMS phenotype. Control rats received daily injections of the selective OTR antagonist d(CH2)5-Tyr(Me)-[Orn8]-vasotocin (dOVT) during postnatal days 2-12, matching the NMS period. At adulthood, behavioral assessments revealed that control+dOVT animals exhibited mechanical and cold thermal hypersensitivity similar to NMS rats, though hot thermal sensitivity was unaffected. Anxiety-like behaviors observed in NMS animals were not reproduced by dOVT treatment. Notably, sex-specific spatial memory deficits emerged: male NMS and female control+dOVT rats showed impaired object location recognition, while females and males in their respective opposite groups remained unaffected. Molecular analyses of spinal cord tissue revealed significant downregulation of GAD65, BDNF, and CD11b in control+dOVT animals. Chloride cotransporters NKCC1 and KCC2 exhibited sexual dimorphism with opposite changes in NMS males versus females and different responses to dOVT. These expressions yet converged on an elevated NKCC1/KCC2 ratio in both sexes, indicating compromised chloride homeostasis despite sex-divergent molecular pathways. These findings demonstrate that developmental OTR dysfunction likely contributes to nociceptive and cognitive consequences of ELS, while anxiety-like phenotypes probably involve additional mechanisms. This work highlights OTR as a critical mediator of neurodevelopmental programming and a potential therapeutic target for mitigating ELS-related disorders.

## Introduction

Adverse experiences such as physical, sexual and emotional abuse during childhood have repeatedly been proven to be detrimental to the psychological and physical development of a child, leading to multiple consequences at adulthood (Felitti et al., 1998; Pechtel and Pizzagalli, 2011; Herzog and Schmahl, 2018). In human, these early life stresses (ELS) and trauma have notably been associated with an increased likelihood of developing several neuropsychiatric disorders (including mood and anxiety disorders), vulnerability to cognitive and emotional dysfunction, as well as shortened lifespan (Felitti et al., 1998; Ladd et al., 2000; Bandelow et al., 2005; Pechtel and Pizzagalli, 2011; Herzog and Schmahl, 2018; Melchior et al., 2022; Waters and Gould, 2022). In order to study more precisely the consequences of these adverse early life conditions, rodent models such as neonatal maternal separation (NMS) have been extensively used. NMS targets a critical period of neurodevelopment (postnatal day 2 (P2) to P12), during which offspring are separated daily from their mothers for 3 hours, inducing chronic neonatal stress. During this specific timespan, essential plastic changes happen in the nervous system, making different brain structures highly sensitive and malleable. At the spinal cord (SC) level, this period is critical for the maturation and organization of spinal cord neurons, critically influenced by neurotrophic factors such as brain-derived neurotrophic factor (BDNF). This growth factor is also a key regulator of the GABAergic system and of chloride homeostasis development, underlying establishment of the nociceptive system (Rivera et al., 1999; Beggs et al., 2002; Stil et al., 2009; Boyce and Mendell, 2014; Verriotis et al., 2016; Melchior et al., 2022). Intense neonatal stress thus induces long-term behavioral consequences such as visceral, mechanical and thermal pain hypersensitivity (Coutinho et al., 2002; Melchior et al., 2022), as well as spinal molecular dysregulation of chloride transporters, neuroinflammatory markers and of oxytocinergic analgesic controls (Melchior et al., 2018; Gazzo et al., 2021). At the supraspinal level, limbic structures such as the amygdala or the hippocampus also undergo intense remodeling during the first postnatal weeks, which makes them primarily susceptible to ELS (Rice and Barone, 2000; McEwen, 2001; Ehrlich et al., 2013; Hoeijmakers et al., 2015; Alves et al., 2022). At adulthood, NMS rodents show neuroendocrine dysregulations and hypothalamo-pituitary-adrenal (HPA) axis hyperactivity, leading to higher anxiety-like behaviors (Plotsky and Meaney, 1993; Ladd et al., 2000; Hoeijmakers et al., 2015), alongside cognitive deficits (Alves et al., 2022; Illouz et al., 2025). In the hippocampus, the GABA excitatory-to-inhibitory switch has been proven to be delayed in NMS animals, inducing disruption of the excitatory/inhibitory (E/I) balance (Furukawa et al., 2017). These changes in basal transmission, associated with altered synaptic plasticity and neuroinflammation have been suggested to drive the mnesic long-term consequences of ELS in a sex-specific way (Aisa et al., 2009; Giridharan et al., 2019; Wang et al., 2020). Indeed, when it comes to learning and memory tasks, females appear to be resilient to NMS effects, whereas both sexes show similar vulnerability to NMS-induced anxiety-like behaviors and nociceptive alterations (Alves et al., 2022; Illouz et al., 2025). However, most studies still do not include females, leaving these differences poorly understood.

In addition, dysregulation of the oxytocinergic system in the hippocampus following NMS has also been highlighted (Lukas et al., 2010; Baracz et al., 2020). However, whether alteration of OTR signaling during the critical early postnatal period is sufficient to trigger NMS-like long-term consequences remains unknown. The nonapeptide Oxytocin (OT) is synthesized by hypothalamic neurons and act as a neurohormone or neurotransmitter depending on whether it is released into the blood through the posterior pituitary gland or into spinal or supraspinal brain structures. This peptide acts preferentially via its G protein-coupled OT receptors (OTR), widely represented at the spinal and supraspinal level (Elands et al., 1988; Tribollet et al., 1992; Mitre et al., 2016). OTR is highly conserved between mammalian species, but some residues can vary and different variants of OTR have been suggested to play a role in neurodevelopmental disorders (Gimpl and Fahrenholz, 2001; LoParo and Waldman, 2015). In fact, OT plays multiple roles around birth and in neurodevelopment, as in parturition, milk ejection or in the newborn response to maternal care (Bales and Perkeybile, 2012). During development, OT promotes the maturation of inhibitory neurotransmission through upregulation of KCC2 expression (Leonzino et al., 2016). Furthermore, OTR signaling in hippocampal neurons has been shown to be neuroprotective, shaping circuit development in young offspring (Cilz et al., 2019; Wu et al., 2021). Later in life, OT displays OTR-mediated analgesic effects after administration in the CNS (Yu et al., 2003; Juif and Poisbeau, 2013). In the central amygdala, OT acts on neuronal activity via OTR-expressing astrocytes, inducing fear suppression and decreased anxiety (Wahis et al., 2021). OTRs are also expressed in both excitatory pyramidal neurons and inhibitory interneurons in the hippocampus, and hippocampal OT modulates neuronal activity, impacting several mnesic and cognitive aspects (Tomizawa et al., 2003; Raam et al., 2017; Cilz et al., 2019; Zagrean et al., 2022).

NMS in male rats was shown to be associated with changes in OTR-mRNA expression and binding sites in several brain regions, suggesting high vulnerability of this system to the early environment (Lukas et al., 2010; Demarchi et al., 2023; Illouz et al., 2025). Moreover, this peptide and its receptor appear to be critical regarding long-term nociceptive consequences of NMS. In fact, NMS rats do not display OTR-mediated stress-induced analgesia (Melchior et al., 2018) and intracerebroventricular administration of OT increases the pain threshold in NMS mice (Amini-Khoei et al., 2017a). OT also attenuates the NMS-induced anxiety and depressive-like behaviors through improving mitochondrial function and decreasing neuroinflammation in the hippocampus (Amini-Khoei et al., 2017b; Najafabadi et al., 2026). In humans, ELS has also been associated with low OT concentrations in cerebrospinal fluid (Heim et al., 2009).

Despite this accumulating evidence that NMS perturbs oxytocinergic signaling, the specific contribution of OTR dysfunction to long-term NMS-induced alterations remains unclear. As mentioned above, OT and OTR play critical roles during early postnatal neurodevelopment, leading us to hypothesize that developmental OTR blockade in CTRL animals would induce key behavioral and molecular consequences of the NMS phenotype. Thus, we administered daily neonatal doses of the selective OTR antagonist dOVT to pups of the control (CTRL) group during the critical P2-P12 period. As this time matches the NMS-period, it allows to isolate the specific contribution of OTR dysfunction during this critical developmental phase. At adulthood, long-term nociceptive, anxiety-like and mnesic outcomes in adult males and females across the three groups (CTRL, CTRL+dOVT and NMS) were assessed. Molecular analyses focused on spinal cord samples, as this structure integrates both nociceptive processing and descending modulatory influences from supraspinal regions, and displays well-characterized molecular alterations following NMS (Melchior et al., 2018; Gazzo et al., 2021). We examined neurotrophic (BDNF), GABAergic (GAD65, NKCC1, KCC2), neuroinflammatory (CD11b, GFAP, TNFα, IL1β), and oxytocinergic (OT, OTR) markers to determine whether developmental OTR blockade induces a similar spinal molecular signature.

## Materials and Methods

### Animals

Pregnant female Sprague-Dawley rats (Charles River, Saint-Germain Nuelles, France) were used in this study and carefully monitored for delivery. The day of birth was referred to as postnatal day 0 (P0). Mother and pups were housed in a temperature (22 ± 2 °C) and humidity (45 ± 10 %) controlled room, under a 12-hour light–dark cycle (lights on at 7:00 am), with ad libitum access to food and tap water. Pups were weaned at P21 and housed in collective cages according to sex. Males and females were used in this study, and results were pooled when no sex-specific difference was observed. Different litters were used to perform behavioral tests (3 litters/group) and molecular analyses (2 litters/group). All procedures were conducted in accordance with EU regulations and approved by the regional ethical committee (CREMEAS authorization numbers APAFIS #41332-2023021317122829 v6).

### Neonatal Maternal Separation

Litters were randomized at birth into two groups: nonseparated control litters and NMS. From postnatal days 2 (P2) to 12 (P12), the litters assigned to the NMS group were taken out of the nest cages three hours a day and put in a different cage under a heating lamp (28 ± 2°C wavelengths between 1400 nm and 3000 nm, one cage/litter). During the whole NMS time, the litters assigned to the control group stayed in their home cages with their mothers and did not get any additional treatment aside from changing their cage bedding, and vehicle (veh) injections. The pups were weaned at P21 and kept in cages with four rats each.

### Drug Treatments

All molecules used in this study were injected intraperitoneally (i.p. 10 µl, 30-gauge needle) daily during the NMS period, i.e., from P2 to P12 (11 injections in total). The injections were performed at the beginning of the separation process. NaCl (0.9 %) was used to control for the effect of daily injections into pups in the CTRL and NMS groups. d(CH2)5-Tyr(Me)-[Orn8]-vasotocin (dOVT, selective oxytocin receptor antagonist, Bachem, Weil am Rhein, Germany) was aliquoted upon receipt at 0.5 mg/mL, then diluted in NaCl 5 minutes before the experiment to 50 µg/kg; and injected CTRL offsprings.

### Behavioral Testing

Rats were tested between P35 and P60. Experiments were performed during the light phase, between 9:00a.m. and 5:00p.m. Prior to behavioral testing, animals were habituated to the experimenter (3 days of handling). In order to eliminate olfactory stimuli, apparatus were always cleaned after testing each animal.

#### Nociception

Mechanical nociception was assessed on P35 to P43 rats with a calibrated forceps (Bioseb, Vitrolles, France). The habituated rat was loosely restrained under a towel masking the eyes, limiting stress by environmental stimulations. The tips of the forceps were placed at each side of the paw, and a gradually increasing force was applied. The pressure producing withdrawal of the paw was noted and corresponded to the nociceptive threshold value.

Thermal hot and cold nociception was measured using hot- and cold-plate tests (Hot-Cold Plate, Bioseb, France). Without habituation, animals were placed in a Plexiglas compartment on a hot or cold plate, maintained at 52°C or 0°C respectively. The latency of onset of the first nociceptive behavior (licking or withdrawal of one or more paws, hobbling, or jumping) was recorded and corresponds to the rat’s thermal nociceptive threshold. If no nociceptive behavior is observed at the end of the first 60 seconds, the animal is removed from the chamber to avoid potential tissue damage.

#### Anxiety-like Behavior

The light-dark box allowing measurement of anxiety-like behavior consisted of two interconnected compartments made of two Plexiglas cylinders (height: 25.5 cm, diameter: 20 cm), one of which was dark and the other exposed to light (170 lux). Rats were placed in the device, and exploration was measured over 5 minutes. The time spent in the dark and light compartments was recorded.

#### Short-term memory

The object recognition and location tests were adapted from the literature (Ennaceur and Delacour, 1988). The open field (OF) consisted of a black wooden arena measuring 65 cm by 65 cm by 45 cm (30 lux). The rats were habituated to the OF for 15 minutes on the 3 days before the trial. In the first part of the novel object recognition test (NORT), two similar objects (2 rectangular plastic boxes, 40cm x 15cm x 5cm) were put inside the chamber at equal distances (10 cm) from the sides. Prior to testing, all objects were assessed for baseline preferences to ensure no inherent biases existed between objects. Animals were placed with their nose facing the wall in a corner of the open field and allowed to explore for 15 minutes. Exploration of the object was considered as soon as the rat had its head oriented toward the object with its nose within 2 cm of the object. One hour later, a second trial took place, in which one object was replaced by a different one (1 round iron box, Ø = 5 cm, h = 35 cm), and exploration was scored for 4 minutes. To avoid preference for one of the objects, the order of the objects was balanced between the testing animals.

The same scheme was used for the object location test (OLT, 3 days habituation sessions, one training phase and a 1-hour testing trial), now using two identical objects but changing their location (2 glass bottles, Ø = 10 cm, h = 30 cm). Rats were always placed with their nose facing the wall, like during the training phase. One hour after training, one of the objects was relocated in the diagonal corner relative to the other object. In the same way as for NORT, which of the two objects (the left or right object) was relocated, as well as the wall upon which it was organized, was counterbalanced.

Each test phase (habituation, training phase, and testing trial) was filmed to be analyzed later, and exploration of the two objects was scored for 4 minutes. Results were expressed as the difference in the exploration time of the two objects divided by the total time spent exploring the objects (discrimination ratio) (Ennaceur and Delacour, 1988; Mendez et al., 2015).

### Molecular Analysis

#### Tissue collection

The lumbar segment of the SC was collected at P55 with n = 10/group, harvested and stored at -80°C.

#### RNA extraction and RT-qPCR

Total RNA was extracted using a protocol adapted from the original procedure of Chomczynski & Sacchi (1987), consisting in 2 independent total RNA extractions separated by a DNAse I treatment (TURBOTM DNase; Ambion, Life technologies, Saint Aubin, France), as previously described in detail (Lelievre et al., 2002). 800 ng RNA were reverse transcripted with the RT iScript kit (Bio-Rad, Marnes-la-Coquette, France). Quantitative PCR was performed using SYBR Green Supermix (Bio-Rad), on the iQ5 Real Time PCR System (Bio-Rad). Amplifications were carried out in 45 cycles (20 s at 95°C, 20 s at 60°C, and 20 s at 72°C). Primer sets for all genes of interest were designed using Oligo6.0 and M fold software and previously used in (Gazzo et al., 2021; Illouz et al., 2025) (primer sequences in Supplementary table 1). Samples were accurately dispensed in duplicates using a robotic workstation (Freedom EVO100; Tecan, Lyon, France), and amplification efficacy given by standard curves was always close to 100 % (± 2 %), while amplification specificity was assessed by a melting curve study. Data were normalized to the housekeeping gene glyceraldehyde 3-phosphate dehydrogenase (GAPDH) since its transcripts remained highly stable among the different samples.

### Statistical Analysis

For each experiment, normality of residuals and homoscedasticity was verified with Shapiro-Wilk and Spearman’s tests prior to performing parametric analyses. Differences were considered statistically significant at p < 0.05. Data are expressed as mean ± standard error of the mean (SEM).

Brown-Forsythe ANOVA followed by Dunnett’s T3 multiple comparisons test was used to compare mechanical nociception between CTRL+veh, NMS+veh and CTRL+dOVT groups, while 1wANOVA followed by Tukey’s multiple comparison test was used concerning thermal hot nociception and anxiety analysis, and Kruskal-Wallis test followed by Dunn’s multiple comparisons test concerning thermal cold nociception. Concerning memory tests, the novel object recognition and object location tasks were analyzed with one sample t-test with 0 as theoretical mean, 0 being the value for which the animals have no preference for any of the objects. Inter- and intra-group effects were assessed using 1- or 2-way ANOVA followed by Tukey’s multiple comparison post-hoc test. Statistical analysis was performed using GraphPad Prism 9 software (La Jolla, USA).

All PCR results were processed using the ΔΔct method (Livak and Schmittgen, 2001) (R studio, version 4.5.1; readxl, dplyr, writexl). Intergroup comparisons of transcript expressions were done using 1wANOVA followed by Tukey’s multiple comparison post-hoc test (GraphPad Prism 9 software, La Jolla, USA).

## Results

Behavioral long-term effects of NMS and dOVT injections were characterized between P35 and P55 (fig.1A). First, mechanical and thermal nociceptive thresholds were assessed (fig.1B), then anxiety in the light-dark box (fig.1C) and short-term recognition and spatial memory (fig.1D).

**Figure 1:**
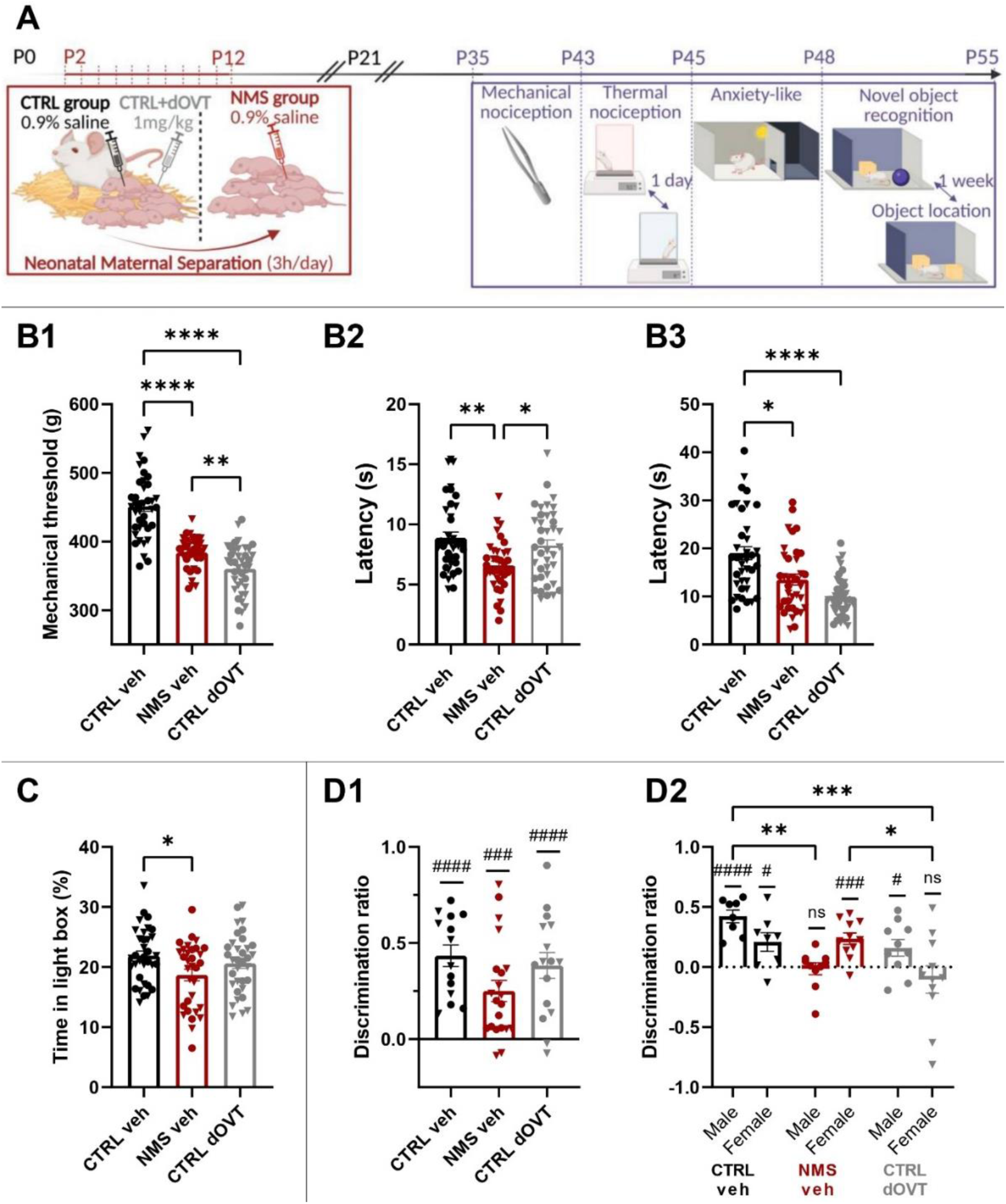
NMS and OTR blockade produce overlapping adult pain phenotypes with distinct affective profiles. **A:** After birth (P0), randomly selected litters underwent daily injection of vehicle or dOVT before undergoing NMS or not. All pups remained with their mothers until weaning at P21. Behavioral testing was conducted between P35 and P55. **B1:** NMS and early dOVT injections induced a decrease in mechanical nociceptive thresholds as evaluated with calibrated forceps. **B2:** Hot plate (52°C) test revealed thermal hypersensitivity in NMS+veh rats, but not in CTRL+dOVT rats. **B3:** Cold plate (0°C) tests highlighted cold hypersensitivity in NMS+veh and CTRL+dOVT rats. **C:** NMS also induced a decrease in time spent in the light compartment in a light-dark box test, which is not the case concerning CTRL+dOVT rats. **D1:** CTRL+veh, NMS+veh and CTL+dOVT rats presented an intact recognition memory. **D2:** NMS+veh rats showed a male-specific (•) spatial memory deficit, and CTRL+dOVT rats showed a female-specific (6) spatial memory deficit. When no differences were observed between male and female rats, statistical significance was assessed with Brown-Forsythe ANOVA followed by Dunnett’s T3 multiple comparisons test (B1), 1wANOVA followed by Tukey’s multiple comparison test (B2, C, D1), Kruskal-Wallis test followed by Dunn’s multiple comparisons test (B3). Concerning the OLT, 2wANOVA followed by Tukey’s multiple comparison post-hoc test revealed sex and group differences. Statistical results were illustrated as follows: p<0.05 (*), p < 0.01 (**), p < 0.001 (***) or p < 0.0001 (****). Memory tests were also analyzed with one sample t-test against 0, and illustrated as follows: p<0.05 (#), p < 0.001 (###) or p < 0.0001 (####). Nociceptive and anxiety tests: Male CTRL+veh: n=19, Female CTRL+veh: n=20, Male NMS+veh: n=19, Female NMS+veh: n=17, Male CTRL+dOVT: n=16, Female CTRL+dOVT: n=23. Memory tests: Male CTRL+veh: n=8, Female CTRL+veh: n=8, Male NMS+veh: n=10, Female NMS+veh: n=11, Male CTRL+dOVT: n=8, Female CTRL+dOVT: n=8.

Compared to the CTRL+veh group, NMS+veh and CTRL+dOVT animals exhibited mechanical and thermal hot nociceptive hypersensitivity at young adulthood (fig. 1B). NMS rats displayed significant 15% lower mean pressure thresholds, while CTRL+dOVT rats also present significantly lower (-20% compared to CTRL+veh) mean paw withdrawal pressure (fig1B1; Brown-Forsythe ANOVA test: P<0.0001, F*(2, 92.88)=64.93; Dunnett’s T3 multiple comparisons test: CTRL+veh vs. NMS+veh P<0.0001, CTRL+veh vs. CTRL+dOVT P<0.0001). Similarly, NMS+veh and CTRL+dOVT withdrawal latencies to noxious cold were respectively 29% and 48% significantly lower than CTRL+veh withdrawal latencies to noxious cold, indicating nociceptive hypersensitivity to cold for both groups (fig1B3; Kruskal-Wallis test: P<0.0001; Dunn’s multiple comparisons test: CTRL+veh vs. NMS+veh P=0.0146, CTRL+veh vs. CTRL+dOVT P<0.0001). Concerning nociceptive heat sensitivity, NMS+veh rats displayed reduced mean withdrawal latencies (-26%), while CTRL+dOVT had similar withdrawal latencies as the CTRL+veh group (fig1B2; 1wANOVA test: P=0.0014, F(2,111)=7.013; Tukey’s multiple comparisons test: CTRL+veh vs. NMS+veh P=0.0012, CTRL+veh vs. CTRL+dOVT P=0.5494). In summary, multimodal nociceptive hypersensitivity is observed in young adult NMS+veh rats, and neonatal dOVT injections to CTRL rats induce a similar phenotype to NMS concerning mechanical and cold thermal nociceptive modalities, but not hot thermal nociception.

Analysis of light-dark box exploration revealed that P45 NMS+veh animals are in average 15% more anxious than CTRL+veh rats, as suggested by the reduced time spent in the light compartment of the light-dark box. In comparison, CTRL+dOVT and CTRL+veh rats spent approximately the same amount of time in this compartment (fig1C; 1wANOVA test: P=0.0374, F(2,97)=3.399; Tukey’s multiple comparisons test: CTRL+veh vs. NMS+veh P=0.0293, CTRL+veh vs. CTRL+dOVT P=0.5486), revealing increased anxiety-like symptoms of NMS rats as described in the literature (Plotsky and Meaney, 1993). Based on these results, it is unlikely that neonatal injection of dOVT alters the long-term anxiety state of rats.

The NORT showed that CTRL+veh, NMS+veh and CTRL+dOVT rats were able to discriminate between a previously encountered object and a novel object, as the means discrimination ratio were all significantly higher than identical exploration of the two objects (Fig.1D1; One sample t-test; CTRL+veh: P<0.0001; t=7.662, df=13 – NMS+veh: P=0.002; t=4.517, df=20 – CTRL+dOVT: P<0.0001; t=5.726, df=15). No intergroup differences were observed (1wANOVA, p=0.0836, F(2,48)=2.614). Concerning short-term spatial memory in the OLT, the inter-group comparison revealed a group and interaction effect, as well as different profiles between male and female among groups (2wANOVA, group: F(2,52)=7.162, p=0.0018 – interaction group x sex: F(2,52) = 7.593, p=0.0013 – Tukey’s multiple comparison test, Male:CTRL veh vs. Male:NMS veh: p=0.0029, Male:CTRL veh vs. Female:CTRL dOVT: p=0.0001, Female:NMS veh vs. Female:CTRL dOVT: p=0.014). One sample t-test confirmed that male and female CTRL+veh rats did recognize the moved object, as their means discrimination ratios were positive (Fig.1D2; One sample t-test; CTRL+veh Male: P<0.0001; t=8.028, df=7 – CTRL+veh Female: P=0.0313; t=2.685, df=7). The same applies for female NMS+veh and male CTRL+dOVT (Fig.1D2; One sample t-test; NMS+veh Female: P=0.0005; t=5.002, df=10 – CTRL+dOVT Male: P=0.0467; t=2.304, df=9). However, male NMS+veh and female CTRL+dOVT have difficulties to discriminate between a previously encountered object location and a novel location, since the average of their discrimination ratios is close to zero, and therefore to the odds (Fig.1D2; One sample t-test; NMS+veh Male: P=0.7752; t=0.2943, df=9 – CTRL+dOVT Female: P=0.379; t=0.9204, df=10).

NMS has been shown to be associated with the expression of several neuroinflammatory mediators in the spinal cord, cerebrum and hippocampus of NMS animals (Gazzo et al., 2021; Illouz et al., 2025), as well as dysregulated neurotrophic factors, chloride homeostasis, and ion channel expression (Juif et al., 2016; Gazzo et al., 2021). Thus, we measured the expression of these markers in the SC to establish whether the NMS-similar behavioral phenotype observed in CTRL+dOVT animals presents the same molecular markers as those observed in the NMS model.

Figure 2A recapitulate no differences in mRNA expression between CTRL+veh and NMS+veh groups regarding the neuroinflammatory markers (CD11b, GFAP, TNFα and IL1β, fig. 2A and 2B). The same is true concerning chloride cotransporters NKCC1 and KCC2 and their ratio, the neurotrophic factor BDNF as well as the oxytocinergic system and (fig. 2A, fig. 2C, fig. 2D and fig. 2E). However, transcripts of the GABAergic marker GAD65 seem under-expressed in NMS rats (NMS+veh: One sample Wilcoxon t-test: P=0.0039). Figure 2A also highlight a significantly decreased mRNA expression of CD11b (CTRL+dOVT: One sample Wilcoxon t-test: P=0.0020), TNFα (CTRL+dOVT: One sample Wilcoxon t-test: P=0.0195), GAD65 (CTRL+dOVT: One sample Wilcoxon t-test: P=0.039), NKCC1 (CTRL+dOVT: One sample Wilcoxon t-test: P=0.0020), KCC2 (CTRL+dOVT: One sample Wilcoxon t-test: P=0.0078), BDNF (CTRL+dOVT: One sample Wilcoxon t-test: P=0.0039) and OTR (CTRL+dOVT: One sample Wilcoxon t-test: P=0.0273) in the CTRL+dOVT group compared to the basal expression value (log2(1), i.e. 0).

**Figure 2:**
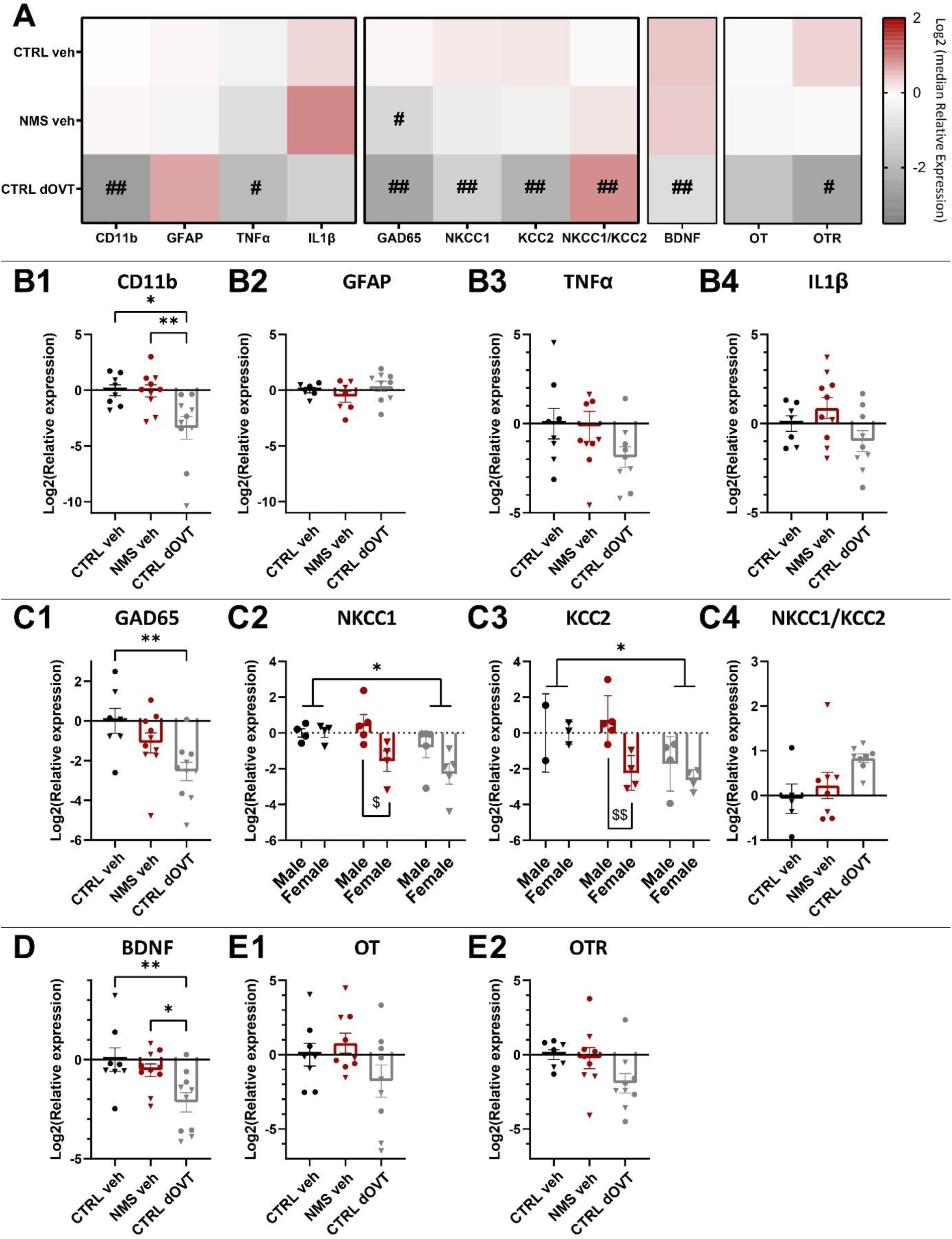
Spinal molecular signature of CTRL+veh, NMS+veh and CTRL+dOVT rats at P55 as measured with RT-qPCR. **A:** CTRL group with early dOVT injections was associated with transcript decreases concerning neuroinflammatory (**B**), inhibition (**C**), neurotrophic (**D**) and oxytocinergic markers (**E**). Median of the results were compared to 0 (theoretical median) with one sample Wilcoxon t-tests, and illustrated as follows: p<0.05 (#), p < 0.01 (##). Statistical significance between groups and sexes was assessed with a 1- or 2wANOVA followed by Tukey’s (group) or Sidak’s (sex) multiple comparison post-hoc test, illustrated as follows: p < 0.05 (*), p < 0.01 (**) (group), p < 0.05 ($), p < 0.01 ($$) (sex). CTRL+veh: n=8 (4 males, 4 females), NMS+veh: n=10 (5 males, 5 females), CTRL+dOVT: n=10 (5 males, 5 females).

In accordance with these results, intergroup comparison point out significant mRNA under-expression in the SC of CTRL+dOVT animals compared to CTRL+veh rats. In fact, figure 2 B1 indicates that spinal CD11b mRNA was 10,5 times less expressed in CTRL+dOVT animals (1wANOVA test: P=0.0041, F(2,25)=6.896; Tukey’s multiple comparisons test: CTRL+veh vs. CTRL+dOVT P=0.0122, NMS+veh vs. CTRL+dOVT P=0.0089). No significant difference was observed for other neuroinflammatory markers (fig. 2B2, 2B3 and 2B4), even though TNFα shows visual underexpression for the CTRL+dOVT group and IL1β presents opposite tendencies between the NMS+veh and CTRL+dOVT groups (1wANOVA test; GFAP: P=0.3136, F(2,19)=1.234 – TNFα: P=0.1814, F (2, 24) = 1.835 – IL1β: P=0.0765, F (2, 23) = 2.88). The relative spinal gene expression of GAD65 was 5.8 fold lower in the SC of CTRL+dOVT rats in comparison to CTRL+veh (fig. 2C1; 1wANOVA test: P=0.0092, F(2,24)=5.729; Tukey’s multiple comparisons test: CTRL+veh vs. CTRL+dOVT P=0.0077). The same is true for NKCC1 and NKCC2 mRNA, which are significantly underexpressed in CTRL+dOVT compared to CTRL+veh (fig. 2C2 and 2C3; 2wANOVA test, main effect of group; NKCC1: P=0.0193; Tukey’s multiple comparisons test: CTRL+veh vs. CTRL+dOVT P=0.0177 – KCC2: P=0.0158; Tukey’s multiple comparisons test: CTRL+veh vs. CTRL+dOVT P=0.0180). Furthermore, a sex-specific difference was observed in the NMS+veh group, with males showing relative expression of NKCC1 and KCC2 opposite to that of dOVT (i.e., overexpression), while mRNA expression in females was similar to that observed in the CTRL+dOVT group (fig. 2C2 and 2C3; 2wANOVA test; NKCC1: main effect of sex: P=0.0095; Sidak’s multiple comparisons test: CTRL+veh vs. CTRL+dOVT P=0.0256 – KCC2: main effect of group: P=0.0158; Tukey’s multiple comparisons test: CTRL+veh vs. CTRL+dOVT P=0.0069). Measurements of NKCC1 and KCC2 transcripts enable the NKCC1/KCC2 ratio to be established in order to assess the efficiency of chloride extrusion in the lumbar spinal cord cells of animals. This ratio NKCC1/KCC2 is higher in NMS+veh and CTRL+dOVT animals, highlighting the weaker inhibitory potential of GABA in these groups (fig. 2C4; Kruskal-Wallis test: P=0.0388). Concerning BDNF, a 4.4 times mRNA underexpression in CTRL+dOVT rats was observed, which was significant (fig. 2D; 1wANOVA test: P=0.0081, F(2,25)=5.866; Tukey’s multiple comparisons test: CTRL+veh vs. CTRL+dOVT P=0.0097, NMS+veh vs. CTRL+dOVT P=0.0432). Figure 2E1 show no differences regarding spinal OT or OTR mRNA expression in NMS+veh and CTRL+dOVT relative to CTRL+veh, even though they tend to be underexpressed in CTRL+dOVT rats (1wANOVA test; OT: P=0.1168, F(2,23)=2.361 – OTR: P=0799, F (2, 22) = 2.84).

## Discussion

The present study demonstrates that selective pharmacological blockade of OTR during the critical P2-P12 period recapitulates key nociceptive and cognitive features of the NMS phenotype, while failing to reproduce anxiogenic components, thereby hinting on OTR-dependent versus OTR-independent mechanisms of early-life programming. Specifically, neonatal dOVT administration induced long-term nociceptive hypersensitivity (mechanical and thermal cold, but not hot) and sex-specific spatial memory deficits (but in a manner opposite to that observed in NMS animals). At the molecular level, CTRL+dOVT animals exhibited significant spinal cord downregulation of GABAergic, neurotrophic, and neuroinflammatory markers. Remarkably, chloride cotransporters NKCC1 and KCC2 displayed pronounced sexual dimorphism, with opposite directional changes in NMS males versus females, and differential magnitude responses to dOVT treatment. Despite these sex-divergent patterns in absolute expression, the NKCC1/KCC2 ratio converged to similar values across both sexes, and to compromised chloride homeostasis in CTRL+dOVT rats. This functional convergence provides a mechanistic explanation for the sexually monomorphic nociceptive phenotype despite underlying molecular heterogeneity. However, unlike NMS animals, dOVT-treated rats did not display increased anxiety-like behaviors, indicating that OTR dysfunction alone does not fully account for the complete NMS phenotype. These findings support the hypothesis that developmental OTR signaling plays a critical mechanistic role in programming long-term nociceptive processing and spatial memory through sex-specific pathways, while suggesting that additional stress-related mechanisms (likely involving HPA axis dysregulation) contribute to the anxiogenic consequences of early life adversity.

The fact that postnatal OTR blockade alone is sufficient to induce lasting mechanical and cold hypersensitivity in rats demonstrates that intact OTR signaling during the neonatal period is essential for the establishment of nociception. Indeed, this developmental window between P2 and P12 coincides with late maturation of nociceptive circuits, including myelination of spinal neurons, as well as essential plastic changes (activity-dependent reorganization of dorsal horn’s fibers) (Fitzgerald, 2005; Melchior et al., 2022). The present article suggests that disruption of neural activity during this critical period, whether through stress or OTR antagonism, alters the refinement and organization of synaptic connectivity (Beggs et al., 2002). OTR-signaling could for example shape the experience-dependent plasticity occurring during development, as suggested in the literature (Zheng et al., 2014). Moreover, GABAergic system and chloride homeostasis gradually mature during the first few weeks after birth in rodents (Ben-Ari, 2002). OTR-signaling notably promotes the maturation of inhibitory neurotransmission through upregulation of KCC2 expression (Leonzino et al., 2016), while also exerting neuroprotective effects during circuit development (Cilz et al., 2019). These findings are consistent with the sustained dysregulation of inhibitory and inflammatory processes observed in CTRL+dOVT rats. At the molecular level, the absence of early OTR signaling would prevent CREB phosphorylation and, consequently, KCC2 transcription (Leonzino et al., 2016). In accordance with this statement, the persistent downregulation observed at P60 could reflect this altered developmental programming. Additionally, the OTR antagonism could induce the decreased BDNF expression observed at P60, which has also been recently suggested in the literature in supraspinal structures (Dayi et al., 2015; Demirtas et al., 2025). This concomitant reduction in BDNF could also alter KCC2 transcription via the BDNF-TrkB pathway (Lee-Hotta et al., 2019).

However, hot thermal sensitivity remained unaffected in CTRL+dOVT animals in contrary to NMS rats. This dissociation suggests that development of thermal hot nociception relies on OTR-independent pathways. The modality-specific effects align with evidence that mechanical, cold, and heat nociception involve partially segregated spinal circuits and molecular mediators (Basbaum et al., 2009). Neonatal injections of dOVT also failed to reproduce the anxiety-like behaviors observed in NMS animals, indicating that lasting anxiety involves alterations beyond developmental OTR dysfunction. In particular, NMS is associated with HPA axis activation and increase in circulating glucocorticoid levels (Plotsky and Meaney, 1993; Ladd et al., 2000), which isn’t recapitulated by selective OTR blockade during the same developmental period. Similarly, the hot thermal hypersensitivity phenotype may involve stress-activated pathways that extend beyond direct OTR signaling blockade. These results indicate that OTR-signaling is a critical mechanism in early stress-induced alterations, but not the only mediator. While OTR blockade is sufficient to program certain nociceptive and cognitive deficits, the full NMS phenotype results from multiple converging pathways.

Spinal cord molecular analyses at P60 point out modest alterations in NMS animals. Trend toward elevated IL1β mRNA emerged, as well as normalization of TNFα transcript, consistent with previous reports at similar timepoints (Gazzo et al., 2021). These subtle changes confirm temporal resolution of inflammatory responses following NMS, and contrast with the pronounced molecular alterations in CTRL+dOVT rats. In both circumstances, early alterations could lead to “developmental miswiring,” i.e., abnormal (early, delayed, arrested, immature, or ectopic) development of neural interactions (Di Martino et al., 2014). This could persist even if molecular markers subsequently normalize, explaining the behavioral alterations dissociated from molecular changes in NMS animals.

Critically, earlier work did not analyze molecular data in a sex-specific manner, while the analysis in this study reveals sexual dimorphism concerning chloride cotransporter regulation in NMS rats. This sex-divergent pattern contrasts with the sex-convergent behavioral nociceptive phenotype, which has already been suggested in the pain literature. In fact, in rodent models of inflammatory and neuropathic pain, males seem to rely on microglial activation and BDNF-mediated KCC2 downregulation, while alternative pathways involving lymphocyte-derived factors (PACAP, leptin) were highlighted in females, with sex-hormones potentially providing protection against BDNF-mediated KCC2 downregulation (Kohno and Tsuda, 2021; Dedek et al., 2022). These mechanistically distinct pathways lead to equivalent mechanical hypersensitivity similarly to developmental programming in this study. However, during development, BDNF promotes KCC2 upregulation and GABAergic maturation (Lee-Hotta et al., 2019), contrasting with its pathological role in adult pain states where excess microglial BDNF drives KCC2 downregulation and sensitization. In our study, neonatal stress resulted in lasting BDNF deficiency in males and females, associated with failure of normal KCC2 upregulation specifically in females. This sex-specific difference in NMS rats as well as the greater magnitude of KCC2 reduction in dOVT-treated females may reflect higher developmental dependence on OTR-signaling in this sex. It is important to note that the NKCC1/KCC2 ratio is preserved despite sex-specific changes in animals subjected to early stress. The coupling of NKCC1 and KCC2 mRNA expression maintains a normalized chloride gradient in the spinal cord of P60 NMS rats, as already demonstrated in the literature at P45 and P100 (Gazzo et al., 2021). Concerning CTRL+dOVT rats, both sexes present an increased NKCC1/KCC2 ratio similar to that of juvenile or adolescent NMS rats (Gazzo et al., 2021), suggesting a decreased GABAergic inhibitory capacity, consistent with equivalent nociceptive hypersensitivity. The divergence between NMS and dOVT effects on chloride cotransporters points out differences in the nature of OTR disruption. In fact, NMS pups are hypothesized to have reduced oxytocinergic signaling during separation period (Melchior et al., 2018), but the oxytocinergic system could still be functional. Conversely, dOVT produces complete OTR-blockade after daily injection, eliminating all endogenous oxytocin signaling on receptors.

Additionally, inverse memory deficits (male-specific in NMS, female-specific in CTRL+dOVT) suggest different supraspinal mechanisms between these two developmental perturbations. In NMS males, spatial memory impairment could emerge, among other mechanisms, because of stress-induced HPA axis hyperactivation with consequent hippocampal glucocorticoid receptors overstimulation (McEwen, 2001; Hoeijmakers et al., 2015). The relative resilience of NMS females confirms previous findings and has been suggested to involve differential development and vulnerability concerning inhibitory and immune systems, as well as neuroprotective effects of sex-hormones (Alves et al., 2022; Illouz et al., 2025). The female-specific vulnerability to dOVT regarding spatial memory also coincides with recent literature (Shehadeh et al., 2025) and may reflect greater developmental dependence on OTR signaling in female hippocampus. In fact, the oxytocinergic system exhibits sexual dimorphism, with higher OT expression in females, but higher OTR expression in males (Dumais and Veenema, 2016). These results raise the possibility that females rely more heavily on OTR-mediated neuroprotection during hippocampal circuit formation than males (Cilz et al., 2019; Wu et al., 2021). This inversion also points out that sex-specific compensatory mechanisms are recruited differently depending on whether the developmental intervention is systemic (NMS, diffuse neuroendocrine responses) versus receptor-specific (dOVT, selective molecular deficit).

In a translational manner, these findings have several implications for understanding early-life adversity in humans. Genetic variations in the OTR gene have been associated with vulnerability to emotional trauma and lower hippocampal volumes (Malhi et al., 2020), as well as increased prevalence to autism spectrum disorder and schizophrenia (LoParo and Waldman, 2015; Veras et al., 2018; Alfimova et al., 2023). Results of the present study corroborates the literature and suggest that individuals carrying low-affinity OTR variants may be predisposed to long-term consequences of childhood trauma, as reduced OTR function during critical developmental periods could compromise nociceptive and cognitive circuit maturation (Gouin et al., 2017; Veras et al., 2018; Alfimova et al., 2023). Importantly, these findings also suggest that different developmental perturbations may produce similar pain phenotypes through distinct mechanisms. Clinically, this underscores the need for personalized approaches that consider not only current symptomatology but also developmental history and potential underlying molecular substrates. The sex-specific vulnerabilities identified in these results further emphasize the need for sex-tailored approaches in both research and clinical contexts.

To conclude, while developmental OTR blockade is sufficient to program long-term nociceptive hypersensitivity and sex-specific cognitive deficits through disrupted GABAergic maturation and chloride homeostasis, the anxiogenic component of NMS requires additional stress-activated pathways, likely involving sustained HPA axis dysregulation. The sexual dimorphism in chloride transporter regulation also reveals sex-specific developmental mechanisms. These findings establish OTR as a critical mediator of neurodevelopmental programming and suggest that OTR-based interventions during critical windows may offer opportunities to mitigate long-term consequences of neonatal adversity. Future investigations should examine hippocampal and amygdalar tissues to identify supraspinal substrates underlying anxiety-like and cognitive alterations following developmental OTR blockade. Mechanistic studies employing pharmacological rescue experiments would test causal relationships between specific molecular alterations and behavioral phenotypes.

## Acknowledgments

Research work has been financed by recurrent funding from the CNRS and the University of Strasbourg, as well as by grants from the Agence Nationale de la Recherche (ANR) as part of the Programme d’Investissement d’Avenir (ANR-17-EURE-022 contract, EURIDOL), the Région Grand Est (ClueDOL, Fonds de Coopération Régionale et de Recherche). PP is a senior member of the Institut Universitaire de France. HI received a fellowship from the french Ministère chargé de l’enseignement supérieur et de la recherche.

## References

Aisa B, Elizalde N, Tordera R, Lasheras B, Del RÃ-o J, RamÃ-rez MJ (2009) Effects of neonatal stress on markers of synaptic plasticity in the hippocampus: Implications for spatial memory. Hippocampus 19:1222–1231.

Alfimova MV, Mikhailova VA, Gabaeva MV, Plakunova VV, Lezheiko TV, Golimbet VE (2023) [Effects of oxytocin pathway gene polymorphisms and adverse childhood experiences on emotion recognition in schizophrenia spectrum disorders]. Zh Nevrol Psikhiatr Im S S Korsakova 123:90–95.

Alves J, de Sá Couto-Pereira N, de Lima RMS, Quillfeldt JA, Dalmaz C (2022) Effects of Early Life Adversities upon Memory Processes and Cognition in Rodent Models. Neuroscience 497:282–307.

Amini-Khoei H, Amiri S, Mohammadi-Asl A, Alijanpour S, Poursaman S, Haj-Mirzaian A, Rastegar M, Mesdaghinia A, Banafshe HR, Sadeghi E, Samiei E, Mehr SE, Dehpour AR (2017a) Experiencing neonatal maternal separation increased pain sensitivity in adult male mice: Involvement of oxytocinergic system. Neuropeptides 61:77–85.

Amini-Khoei H, Mohammadi-Asl A, Amiri S, Hosseini M-J, Momeny M, Hassanipour M, Rastegar M, Haj-Mirzaian A, Mirzaian AH-, Sanjarimoghaddam H, Mehr SE, Dehpour AR (2017b) Oxytocin mitigated the depressive-like behaviors of maternal separation stress through modulating mitochondrial function and neuroinflammation. Prog Neuropsychopharmacol Biol Psychiatry 76:169–178.

Bales KL, Perkeybile AM (2012) Developmental experiences and the oxytocin receptor system. Horm Behav 61:313–319.

Bandelow B, Krause J, Wedekind D, Broocks A, Hajak G, Rüther E (2005) Early traumatic life events, parental attitudes, family history, and birth risk factors in patients with borderline personality disorder and healthy controls. Psychiatry Res 134:169–179.

Baracz SJ, Everett NA, Cornish JL (2020) The impact of early life stress on the central oxytocin system and susceptibility for drug addiction: Applicability of oxytocin as a pharmacotherapy. Neurosci Biobehav Rev 110:114–132.

Basbaum AI, Bautista DM, Scherrer G, Julius D (2009) Cellular and Molecular Mechanisms of Pain. Cell 139:267–284.

Beggs S, Torsney C, Drew LJ, Fitzgerald M (2002) The postnatal reorganization of primary afferent input and dorsal horn cell receptive fields in the rat spinal cord is an activity-dependent process. Eur J Neurosci 16:1249–1258.

Ben-Ari Y (2002) Excitatory actions of gaba during development: the nature of the nurture. Nat Rev Neurosci 3:728–739.

Boyce VS, Mendell LM (2014) Neurotrophins and spinal circuit function. Front Neural Circuits 8 Available at: https://www.frontiersin.org/journals/neural-circuits/articles/10.3389/fncir.2014.00059/full [Accessed September 22, 2025].

Chomczynski P, Sacchi N (1987) Single-step method of RNA isolation by acid guanidinium thiocyanate-phenol-chloroform extraction. Anal Biochem 162:156–159.

Cilz NI, Cymerblit-Sabba A, Young WS (2019) Oxytocin and vasopressin in the rodent hippocampus. Genes Brain Behav 18:e12535.

Coutinho SV, Plotsky PM, Sablad M, Miller JC, Zhou H, Bayati AI, McRoberts JA, Mayer EA (2002) Neonatal maternal separation alters stress-induced responses to viscerosomatic nociceptive stimuli in rat. Am J Physiol-Gastrointest Liver Physiol 282:G307–G316.

Dayi A, Cetin F, Sisman AR, Aksu I, Tas A, Gönenc S, Uysal N (2015) The Effects of Oxytocin on Cognitive Defect Caused by Chronic Restraint Stress Applied to Adolescent Rats and on Hippocampal VEGF and BDNF Levels. Med Sci Monit 21:69–75.

Dedek A, Xu J, Lorenzo L-É, Godin AG, Kandegedara CM, Glavina G, Landrigan JA, Lombroso PJ, De Koninck Y, Tsai EC, Hildebrand ME (2022) Sexual dimorphism in a neuronal mechanism of spinal hyperexcitability across rodent and human models of pathological pain. Brain 145:1124–1138.

Demarchi L, Sanson A, Bosch OJ (2023) Brief versus long maternal separation in lactating rats: Consequences on maternal behavior, emotionality, and brain oxytocin receptor binding. J Neuroendocrinol 35:e13252.

Demirtas H, Acikgoz B, Arslankiran A, Dalkiran B, Kiray A, Kiray M, Dayi A, Aksu I (2025) Effects of oxytocin on behavior and neurotrophic factors in the brain of aged female rats exposed to chronic social isolation. Neuropeptides 112:102532.

Di Martino A, Fair DA, Kelly C, Satterthwaite TD, Castellanos FX, Thomason ME, Craddock RC, Luna B, Leventhal BL, Zuo X-N, Milham MP (2014) Unraveling the Miswired Connectome: A Developmental Perspective. Neuron 83:1335–1353.

Dumais KM, Veenema AH (2016) Vasopressin and oxytocin receptor systems in the brain: Sex differences and sex-specific regulation of social behavior. Front Neuroendocrinol 40:1–23.

Ehrlich DE, Ryan SJ, Hazra R, Guo J-D, Rainnie DG (2013) Postnatal maturation of GABAergic transmission in the rat basolateral amygdala. J Neurophysiol 110:926–941.

Elands J, Barberis C, Jard S (1988) [3H]-[Thr4,Gly7]OT: a highly selective ligand for central and peripheral OT receptors. Am J Physiol-Endocrinol Metab 254:E31–E38.

Ennaceur A, Delacour J (1988) A new one-trial test for neurobiological studies of memory in rats. 1: Behavioral data. Behav Brain Res 31:47–59.

Felitti VJ, Anda RF, Nordenberg D, Williamson DF, Spitz AM, Edwards V, Koss MP, Marks JS (1998) Relationship of Childhood Abuse and Household Dysfunction to Many of the Leading Causes of Death in Adults: The Adverse Childhood Experiences (ACE) Study. Am J Prev Med 14:245–258.

Fitzgerald M (2005) The development of nociceptive circuits. Nat Rev Neurosci 6:507–520.

Furukawa M, Tsukahara T, Tomita K, Iwai H, Sonomura T, Miyawaki S, Sato T (2017) Neonatal maternal separation delays the GABA excitatory-to-inhibitory functional switch by inhibiting KCC2 expression. Biochem Biophys Res Commun 493:1243–1249.

Gazzo G, Melchior M, Caussaint A, Gieré C, Lelièvre V, Poisbeau P (2021) Overexpression of chloride importer NKCC1 contributes to the sensory-affective and sociability phenotype of rats following neonatal maternal separation. Brain Behav Immun 92:193–202.

Gimpl G, Fahrenholz F (2001) The Oxytocin Receptor System: Structure, Function, and Regulation. Physiol Rev Available at: https://journals.physiology.org/doi/10.1152/physrev.2001.81.2.629 [Accessed May 22, 2024].

Giridharan VV, Réus GZ, Selvaraj S, Scaini G, Barichello T, Quevedo J (2019) Maternal deprivation increases microglial activation and neuroinflammatory markers in the prefrontal cortex and hippocampus of infant rats. J Psychiatr Res 115:13–20.

Gouin JP, Zhou QQ, Booij L, Boivin M, Côté SM, Hébert M, Ouellet-Morin I, Szyf M, Tremblay RE, Turecki G, Vitaro F (2017) Associations among oxytocin receptor gene (OXTR) DNA methylation in adulthood, exposure to early life adversity, and childhood trajectories of anxiousness. Sci Rep 7.

Heim C, Young LJ, Newport DJ, Mletzko T, Miller AH, Nemeroff CB (2009) Lower CSF oxytocin concentrations in women with a history of childhood abuse. Mol Psychiatry 14:954–958.

Herzog JI, Schmahl C (2018) Adverse Childhood Experiences and the Consequences on Neurobiological, Psychosocial, and Somatic Conditions Across the Lifespan. Front Psychiatry 9 Available at: https://www.frontiersin.org/articles/10.3389/fpsyt.2018.00420 [Accessed November 21, 2023].

Hoeijmakers L, Lucassen PJ, Korosi A (2015) The interplay of early-life stress, nutrition, and immune activation programs adult hippocampal structure and function. Front Mol Neurosci 7 Available at: https://www.frontiersin.org/articles/10.3389/fnmol.2014.00103 [Accessed March 27, 2024].

Illouz H, Menger Y, Schaack O, Lelievre V, Poisbeau P (2025) Sex-specific spatial memory deficits associated with region-specific neuroinflammatory changes in the dorsal hippocampus of rats exposed to neonatal repeated maternal separation. Brain Behav Immun 129:388–398.

Juif P-E, Poisbeau P (2013) Neurohormonal effects of oxytocin and vasopressin receptor agonists on spinal pain processing in male rats. PAIN® 154:1449–1456.

Kohno K, Tsuda M (2021) Role of microglia and P2X4 receptors in chronic pain. PAIN Rep 6:e864.

Ladd CO, Huot RL, Thrivikraman KV, Nemeroff CB, Meaney MJ, Plotsky PM (2000) Chapter 7 - Long-term behavioral and neuroendocrine adaptations to adverse early experience. In: Progress in Brain Research (Mayer EA, Saper CB, eds), pp 81–103 The Biological Basis for Mind Body Interactions. Elsevier. Available at: https://www.sciencedirect.com/science/article/pii/S0079612308621329 [Accessed August 26, 2024].

Lee-Hotta S, Uchiyama Y, Kametaka S (2019) Role of the BDNF-TrkB pathway in KCC2 regulation and rehabilitation following neuronal injury: A mini review. Neurochem Int 128:32–38.

Lelievre V, Hu Z, Byun J-Y, Ioffe Y, Waschek JA (2002) Fibroblast growth factor-2 converts PACAP growth action on embryonic hindbrain precursors from stimulation to inhibition. J Neurosci Res 67:566–573.

Leonzino M, Busnelli M, Antonucci F, Verderio C, Mazzanti M, Chini B (2016) The Timing of the Excitatory-to-Inhibitory GABA Switch Is Regulated by the Oxytocin Receptor via KCC2. Cell Rep 15:96–103.

Livak KJ, Schmittgen TD (2001) Analysis of Relative Gene Expression Data Using Real-Time Quantitative PCR and the 2−ΔΔCT Method. Methods 25:402–408.

LoParo D, Waldman ID (2015) The oxytocin receptor gene (OXTR) is associated with autism spectrum disorder: a meta-analysis. Mol Psychiatry 20:640–646.

Lukas M, Bredewold R, Neumann ID, Veenema AH (2010) Maternal separation interferes with developmental changes in brain vasopressin and oxytocin receptor binding in male rats. Neuropharmacology 58:78–87.

Malhi GS, Das P, Outhred T, Dobson-Stone C, Bell E, Gessler D, Bryant R, Mannie Z (2020) Interactions of OXTR rs53576 and emotional trauma on hippocampal volumes and perceived social support in adolescent girls. Psychoneuroendocrinology 115:104635.

McEwen BS (2001) Plasticity of the Hippocampus: Adaptation to Chronic Stress and Allostatic Load. Ann N Y Acad Sci 933:265–277.

Melchior M, Juif P-E, Gazzo G, Petit-Demoulière N, Chavant V, Lacaud A, Goumon Y, Charlet A, Lelièvre V, Poisbeau P (2018) Pharmacological rescue of nociceptive hypersensitivity and oxytocin analgesia impairment in a rat model of neonatal maternal separation. Pain 159:2630–2640.

Melchior M, Kuhn P, Poisbeau P (2022) The burden of early life stress on the nociceptive system development and pain responses. Eur J Neurosci 55:2216–2241.

Mendez M, Arias N, Uceda S, Arias JL (2015) c-Fos expression correlates with performance on novel object and novel place recognition tests. Brain Res Bull 117:16–23.

Mitre M, Marlin BJ, Schiavo JK, Morina E, Norden SE, Hackett TA, Aoki CJ, Chao MV, Froemke RC (2016) A Distributed Network for Social Cognition Enriched for Oxytocin Receptors. J Neurosci 36:2517–2535.

Najafabadi PS, Shamsara A, Mirzaie V, Sheibani V, Ahmadi M, Joushi S, Basiri M (2026) Effects of intranasal oxytocin administration on histophysiology of the hippocampus in maternally separated adolescent male rats. Behav Brain Res 500:115983.

Pechtel P, Pizzagalli DA (2011) Effects of early life stress on cognitive and affective function: an integrated review of human literature. Psychopharmacology (Berl) 214:55–70.

Plotsky PM, Meaney MJ (1993) Early, postnatal experience alters hypothalamic corticotropin-releasing factor (CRF) mRNA, median eminence CRF content and stress-induced release in adult rats. Brain Res Mol Brain Res 18:195–200.

Raam T, McAvoy KM, Besnard A, Veenema AH, Sahay A (2017) Hippocampal oxytocin receptors are necessary for discrimination of social stimuli. Nat Commun 8:2001.

Rice D, Barone S (2000) Critical periods of vulnerability for the developing nervous system: evidence from humans and animal models. Environ Health Perspect 108 Suppl 3:511–533.

Rivera C, Voipio J, Payne JA, Ruusuvuori E, Lahtinen H, Lamsa K, Pirvola U, Saarma M, Kaila K (1999) The K+/Cl- co-transporter KCC2 renders GABA hyperpolarizing during neuronal maturation. Nature 397:251–255.

Shehadeh E, Rajan Narattil N, Kritman M, Maroun M (2025) Sex-dimorphic oxytocin regulation of CA1-dependent spatial memory and synaptic plasticity in juvenile rats. Behav Brain Funct 21:41.

Stil A, Liabeuf S, Jean-Xavier C, Brocard C, Viemari J-C, Vinay L (2009) Developmental up-regulation of the potassium–chloride cotransporter type 2 in the rat lumbar spinal cord. Neuroscience 164:809–821.

Tomizawa K, Iga N, Lu Y-F, Moriwaki A, Matsushita M, Li S-T, Miyamoto O, Itano T, Matsui H (2003) Oxytocin improves long-lasting spatial memory during motherhood through MAP kinase cascade. Nat Neurosci 6:384–390.

Tribollet E, Dubois-Dauphin M, Dreifuss JJ, Barberis C, Jard S (1992) Oxytocin Receptors in the Central Nervous System. Ann N Y Acad Sci 652:29–38.

Veras AB, Getz M, Froemke RC, Nardi AE, Alves GS, Walsh-Messinger J, Chao MV, Kranz TM, Malaspina D (2018) Rare missense coding variants in oxytocin receptor (OXTR) in schizophrenia cases are associated with early trauma exposure, cognition and emotional processing. J Psychiatr Res 97:58–64.

Verriotis M, Fabrizi L, Lee A, Cooper RJ, Fitzgerald M, Meek J (2016) Mapping Cortical Responses to Somatosensory Stimuli in Human Infants with Simultaneous Near-Infrared Spectroscopy and Event-Related Potential Recording. eNeuro 3:ENEURO.0026-16.2016.

Wahis J et al. (2021) Astrocytes mediate the effect of oxytocin in the central amygdala on neuronal activity and affective states in rodents. Nat Neurosci 24:529–541.

Wang A, Zou X, Wu J, Ma Q, Yuan N, Ding F, Li X, Chen J (2020) Early-Life Stress Alters Synaptic Plasticity and mTOR Signaling: Correlation With Anxiety-Like and Cognition-Related Behavior. Front Genet 11:590068.

Waters RC, Gould E (2022) Early Life Adversity and Neuropsychiatric Disease: Differential Outcomes and Translational Relevance of Rodent Models. Front Syst Neurosci 16 Available at: https://www.frontiersin.org/articles/10.3389/fnsys.2022.860847 [Accessed March 18, 2024].

Wu Z, Xie C, Kuang H, Wu J, Chen X, Liu H, Liu T (2021) Oxytocin mediates neuroprotection against hypoxic-ischemic injury in hippocampal CA1 neuron of neonatal rats. Neuropharmacology 187:108488.

Yu S-Q, Lundeberg T, Yu L-C (2003) Involvement of oxytocin in spinal antinociception in rats with inflammation. Brain Res 983:13–22.

Zagrean A-M, Georgescu I-A, Iesanu MI, Ionescu R-B, Haret RM, Panaitescu AM, Zagrean L (2022) Oxytocin and vasopressin in the hippocampus. Vitam Horm 118:83–127.

Zheng J-J, Li S-J, Zhang X-D, Miao W-Y, Zhang D, Yao H, Yu X (2014) Oxytocin mediates early experience–dependent cross-modal plasticity in the sensory cortices. Nat Neurosci 17:391–399.

