## Supplemental Table 1 for "Oxytocin receptor dysfunction during critical neurodevelopment programs lasting pain hypersensitivity and sex-specific cognitive deficits"

Supplementary Table 1: Primer sequences used in this study.

| **Gene** | **Primers sequences (forward/reverse)** | **NCBI reference** |
| --- | --- | --- |
| **GAPDH** | GGCCTTCCGTGTTCCTAC  TGTCATCATACTTGGCAGGTT | NM017008 |
| **CD11b** | CTGCCTCAGGGATCCGTAAAG  CCTCTGCCTCAGGAATGACATC | NM012711 |
| **GFAP** | TGGAGAGAATTGAATCGC  CCGGAGTTCTCGAACTTCCTC | NM010277 |
| **TNFα** | GGGCTGTACCTTATCTACTCC  TATGAAATGGCAAATCGG | NM012675 |
| **IL1β** | TCCTCTGTGACTCGTGGGAT  CGAGGCATTTTTGTTGTTCAT | NM031512 |
| **GAD65** | GACCTGTCCTATGACACGG  AGCTTCAAATCCAGTAGTCCC | NM012563.2 |
| **NKCC1** | GGGCCTCCTCACACGAAGAA  TGAGGAGCCGAGGGTACTTCA | NM019229 |
| **KCC2** | ACTACAGCTGGCCACCTCGC  ATGCTGCCCTCAGAGAAACGC | NM134363 |
| **BDNF** | CACAGTCCTGGAGAAAGTCCC  CCTTCCTTCGTGTAACCCAT | NM012513 |
| **OT** | CTGCTTCGGGGCACCG  GCAAGGCTTCTGGCCAGACTG | NM012996 |
| **OTR** | CCCGTGAAGAGCATGTAGATC  CAAAGAAGCTTCTGCCTTCAT | NM012871 |
